# Evidence for immediate enhancement of hippocampal memory encoding by network-targeted theta-burst stimulation during concurrent fMRI

**DOI:** 10.1101/2020.02.19.956466

**Authors:** Molly S. Hermiller, Yu Fen Chen, Todd B. Parrish, Joel L. Voss

## Abstract

The hippocampus supports episodic memory via interaction with a distributed brain network. Previous experiments using network-targeted noninvasive brain stimulation have identified episodic memory enhancements and modulation of activity within the hippocampal network. However, mechanistic insights were limited because these effects were measured long after stimulation and therefore could have reflected various neuroplastic aftereffects with extended timecourses. In this experiment with human subjects of both sexes, we tested for immediate stimulation impact on encoding-related activity of the hippocampus and immediately adjacent medial-temporal cortex by delivering theta-burst transcranial magnetic stimulation (TBS) concurrent with fMRI, as an immediate impact of stimulation would suggest an influence on neural activity. We reasoned that TBS would be particularly effective for influencing the hippocampus because rhythmic neural activity in the theta band is associated with hippocampal memory processing. First, we demonstrated that it is possible to obtain robust fMRI correlates of task-related activity during concurrent TBS. We then identified immediate effects of TBS on encoding of visual scenes. Brief volleys of TBS targeting the hippocampal network increased activity of the targeted (left) hippocampus during scene encoding and increased subsequent recollection. Stimulation did not influence activity during an intermixed numerical task with no memory demand. Control conditions using beta-band and out-of-network stimulation also did not influence hippocampal activity or recollection. TBS targeting the hippocampal network therefore immediately impacted hippocampal memory processing. This suggests direct, beneficial influence of stimulation on hippocampal neural activity related to memory and supports the role of theta-band activity in human episodic memory.

**Significance Statement:** Can noninvasive stimulation directly impact function of indirect, deep-brain targets such as the hippocampus? We tested this by targeting an accessible region of the hippocampal network via transcranial magnetic stimulation during concurrent fMRI. We reasoned that theta-burst stimulation would be particularly effective for impacting hippocampal function, as this stimulation rhythm should resonate with the endogenous theta-nested-gamma activity prominent in hippocampus. Indeed, theta-burst stimulation targeting the hippocampal network immediately impacted hippocampal activity during encoding, improving memory formation as indicated by enhanced later recollection. Rhythm- and location-control stimulation conditions had no such effects. These findings suggest a direct influence of noninvasive stimulation on hippocampal neural activity and highlight that the theta-burst rhythm is relatively privileged in its ability to influence hippocampal memory function.

## Introduction

The hippocampus exhibits theta-band (~4-8 Hz) oscillatory neural activity that is thought to provide a temporal framework for episodic memory (Buzsaki, 2002; Lisman and Jensen, 2013; Herweg et al., 2020). Episodic memory involves hippocampal interaction with a distributed network (Squire and Zola-Morgan, 1991; Eichenbaum et al., 2007; Ranganath and Ritchey, 2012) that exhibits interregional synchrony of memory-related activity preferentially in the theta band (Fell et al., 2001; Buzsaki and Draguhn, 2004; Foster et al., 2013; Lisman and Jensen, 2013; Staudigl and Hanslmayr, 2013). Although the functional significance of hippocampal theta activity has been tested via stimulation in rodents (Shirvalkar et al., 2010; Zutshi et al., 2018), such direct functional tests present major challenges for human experimentation.

Stimulation using a theta-rhythmic pattern, as in theta-burst stimulation (TBS; high-frequency stimulation delivered in a theta rhythm), should be capable of testing the role of theta in episodic memory. This is because TBS mimics the endogenous theta rhythm thought to support hippocampal memory processing and hippocampal network synchronization, and therefore may optimally influence this network’s function via activity entrainment (Buzsaki, 2002; Thut et al., 2011b; Romei et al., 2016). Direct stimulation of the hippocampus is not possible in human subjects without neurosurgery, but noninvasive stimulation can directly test network functional properties (Fox et al., 2012). Multiple experiments using transcranial magnetic stimulation (TMS) have shown that it is possible to influence episodic memory by targeting the hippocampal network (Hebscher and Voss, 2020). In those experiments, network-targeted TMS increased hippocampal network fMRI connectivity, memory-related fMRI activity, and/or improved memory performance for hours to weeks post-stimulation e.g., (Wang et al., 2014; Kim et al., 2018; Tambini et al., 2018; Freedberg et al., 2019; Hermiller et al., 2019b; Warren et al., 2019). Consistent with the hypothesized importance of theta for memory, one study found that network-targeted TBS had greater impact on memory accuracy and memory-related hippocampal fMRI connectivity than did a non-theta (20-Hz) control frequency (Hermiller et al., 2019a). However, a weakness of previous noninvasive stimulation experiments with respect to mechanistic interpretation is that these studies measured long-lasting aftereffects of stimulation (ranging from minutes to weeks), which could be mediated by a variety of indirect neuroplasticity mechanisms (Thickbroom, 2007). Evidence for preferred influence of TBS on memory-related neural activity would require immediate assessment of stimulation impact.

To address this, we delivered TBS to a hippocampal-network-targeted (HNT) location in the parietal cortex during concurrent fMRI while subjects performed a memory task. We developed custom fMRI parameters that allowed TBS as well as control-frequency (12.5 Hz) stimulation during scanning. These scanning parameters were suitable for our focus on the effects of TBS on hippocampus and adjacent cortex of the medial temporal lobe (MTL), as high-quality fMRI signals were obtained from these areas.

Human subjects studied complex visual scenes that were each immediately preceded by either HNT TBS or by one of several control stimulation conditions (Fig. 1). We hypothesized that HNT TBS delivered in the seconds immediately preceding individual stimuli would increase fMRI activity evoked by scenes in the hippocampus, reflecting the effects of successful entrainment of the hippocampal theta rhythm by TBS. Based on robust evidence that increased hippocampal activity during encoding is associated with later recollection (Paller and Wagner, 2002; Kim, 2011; Rugg et al., 2012) and previous findings that hippocampal network-targeted stimulation enhances recollection (Hebscher and Voss, 2020), we predicted increased recollection for scenes encoded with HNT TBS. We included various control conditions, including controls for the cognitive task (memory versus non-memory), stimulation rhythm (TBS versus non-theta), stimulation location (HNT versus out-of-network), and hemisphere (targeted left versus non-targeted right hippocampus). Because the experiment involved a randomized trial-based stimulation design, immediate and selective effects of HNT TBS on hippocampal memory-related activity would indicate that this type of stimulation impacted hippocampal neural activity, rather than longer-lasting neuroplasticity processes that would carry over to affect other trials, and would thereby support the role of theta in human memory formation.

**Figure 1.**
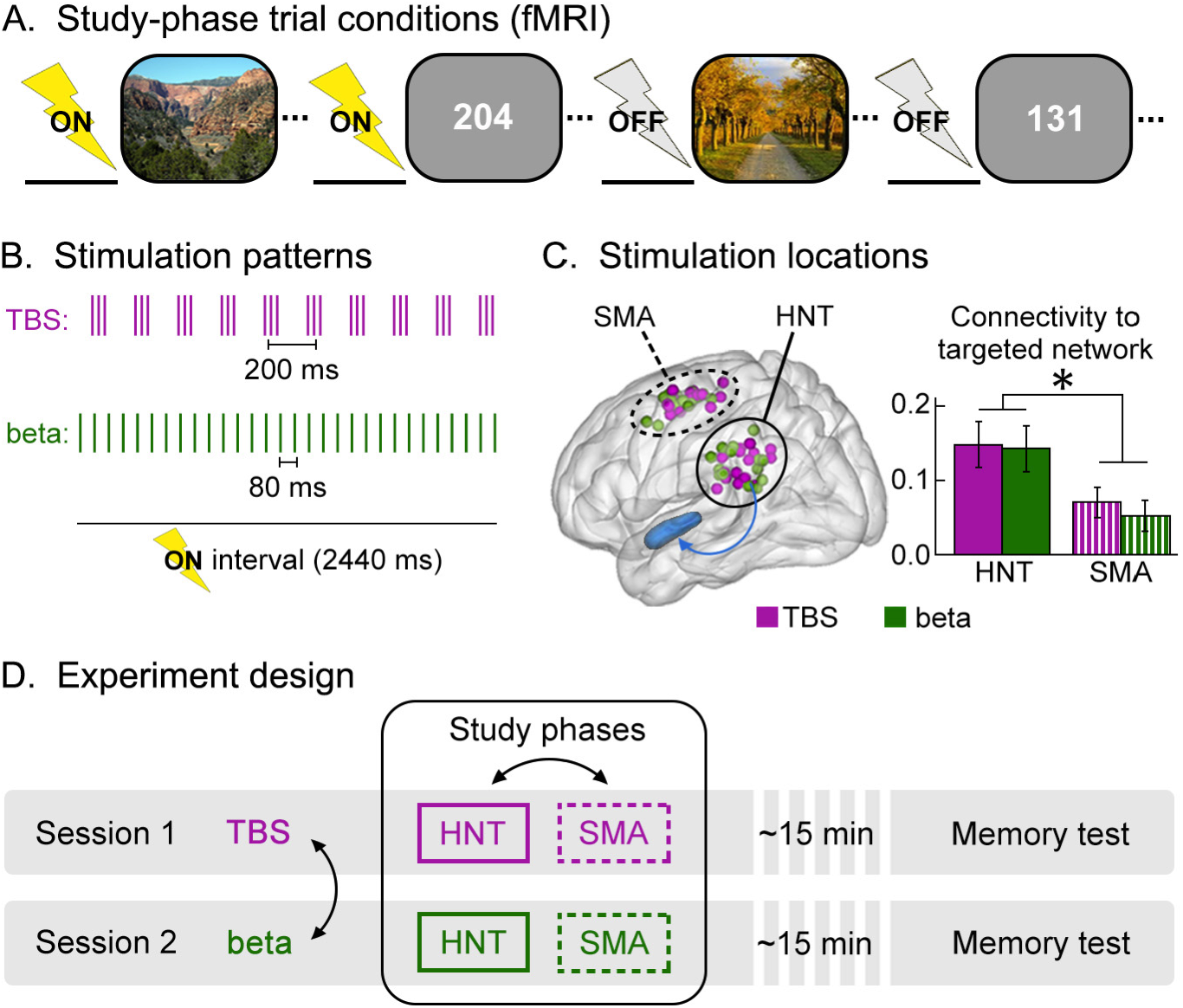
Trial-specific stimulation during episodic memory formation. (**A**) Scene-encoding and numeric-judgment trials were randomly intermixed during each study phase, with ~2 s of stimulation delivered immediately before stimulus onset for a subset of trials (ON) and no preceding stimulation for remaining trials (OFF). Study phases were completed during fMRI scanning with memory test phases after scanning. There were four stimulation conditions for ON and OFF trials. (**B**) Stimulation was delivered as either a theta-burst pattern (TBS: 50 Hz triplet pulses delivered at 5 Hz) or at beta (single pulses delivered at 12.5 Hz). These conditions had the same overall number of pulses during each stimulation period delivered at the same intensity. (**C**) Stimulation was delivered to the HNT parietal location (based on its fMRI connectivity with left hippocampus, as depicted by the blue arrow), or a control out-of-network SMA location. Achieved stimulation locations confirmed via MRI for each condition and subject are indicated by colorized spheres on a template brain. Bar plots represent mean ±s.e.m. baseline resting-state fMRI connectivity of the subject-specific stimulation locations with the hippocampal network, confirming relatively higher connectivity for the HNT than SMA location. *P<0.05 main effect of stimulation location by one-way rmANOVA. (**D**) HNT or SMA locations were targeted for one of the two study phases in each experimental session. After both study phases were complete, subjects exited the scanner for a ~15 min break before taking the memory test. A different stimulation pattern (TBS or beta) was used for each experimental session. Black arrows indicate stimulation conditions with order counterbalanced across subjects.

## Materials and Methods

### Overview

Following a baseline session, subjects completed a two-day experiment in which they attempted to remember complex visual scenes that were each immediately preceded by different stimulation conditions, performed during fMRI scanning. The main condition of interest was TBS delivered to a hippocampal-network targeted (HNT) location in the parietal cortex immediately before the onset of scenes. Several control conditions were used to test specificity. Subjects also received stimulation immediately before the presentation of numeric judgments, interleaved randomly with the scenes throughout the task (Fig. 1A). We expected no effect of stimulation on hippocampal activity for this condition, as hippocampal activity is generally not evoked by numeric judgments (Stark and Squire, 2001) and therefore would not increase via direct effects of stimulation on hippocampal neural activity. Furthermore, the same scene and number conditions were administered in three control stimulation conditions: (i) a different stimulation rhythm (beta; 12.5 Hz) (Fig. 1B) applied to the same HNT location, (ii) TBS applied to a control location in the supplementary motor area (SMA) outside the hippocampal network (Fig. 1C), and (iii) beta stimulation of the SMA location. None of these control conditions were expected to influence hippocampal activity. Finally, stimulation was not delivered for a subset of scene and number trials, providing a no-stimulation (“off”) control condition. Scene and number trials with and without stimulation were intermixed throughout scanning sessions, guarding against confounding influences such as stimulation-induced fMRI artifact and stimulation carry-over effects across trials. All conditions were administered in each subject using a within-subjects counterbalanced design over two experimental sessions (Fig. 1D). As indicated above, our *a priori* hypotheses were that the HNT TBS encoding condition would be unique among all stimulation conditions in producing: (1) increased likelihood of later recollection during the memory test, and (2) increased activity in the left (targeted) hippocampus during encoding of scenes that would be later recollected during the memory test.

### Subjects

Adult human subjects passed standard MRI and TMS safety screenings (Rossi et al., 2009), reported no present use of psychoactive drugs, and were free of known neurological and psychiatric conditions. Datasets from 16 subjects were included in all reported analyses (8 females, 8 males; ages 20-35 years, average age=27.6, SD=4.3). Data from two additional subjects were collected but excluded from all reported analyses due to poor behavioral performance (overall miss rate > 50%). In addition, data collection was attempted from three additional subjects but failed due to technical malfunction (n=2) or attrition (n=1). Subjects gave written informed consent approved by the Northwestern University Institutional Review Board and were paid for participation. The sample size of N=16 was chosen to match or exceed previous experiments that demonstrated memory improvement for stimuli encoded following short volleys of TMS (i.e., <2 s TMS immediately before stimulus onset) (Kohler et al., 2004; Demeter et al., 2016).

### Baseline session

Subjects completed a baseline session to determine stimulation locations and intensity prior to two experimental sessions, performed on different days (described below).

#### Baseline MRI to determine stimulation locations

Resting-state fMRI and structural MRI were collected using a 3T Siemens PRISMA scanner with a 64-channel head/neck coil. Baseline resting-state functional images were acquired using a bloodoxygenation-level-dependent (BOLD) contrast sensitive gradient-echo echo-planar imaging (EPI) pulse sequence (270 frames; TE 20 ms; TR 2000.0 ms; flip angle 80°; voxel resolution 1.7 mm isotropic; 70 ascending axial slices; 210×203 mm FOV; scan duration 9 min). During the resting-state scan subjects were instructed to lie as still as possible, to keep their eyes open and focused on a fixation cross presented in the center of the screen, and to let their minds wander. Structural images were acquired using a T1-weighted MPRAGE sequence (176 frames; TE 1.69 ms; TR 2170 ms; TI 1100 ms; flip angle 7°; voxel resolution 1.0 mm isotropic; 256×256 mm FOV; GRAPPA acceleration of a factor of 2; scan duration 6.36 min).

Baseline scans were submitted to resting-state fMRI connectivity analysis to determine stimulation locations. All fMRI analyses used AFNI (Cox, 1996) and were visualized with the BrainNet Viewer Matlab (The MathWorks, Inc. Natick, MA, USA) toolbox (Xia et al., 2013) on a smoothed Colin27 template. Anatomical scans were skull-stripped *(3dSkullStrip)* and co-registered to standardized space using the Colin27 template *(auto_tlrc).* Preprocessing of the functional volumes included outlier suppression *(3dDespike),* slice timing and motion correction *(3dvolreg),* and co-registration to the anatomical scan *(align_epi_anat).* The transformations were applied simultaneously in a single resampling step *(3dAllineate).* Motion parameters were calculated for each volume as the Euclidean norm of the first difference of six motion estimates (three translation and three rotation). Volumes with excessive motion (>0.2 mm), as well as the previous volume, were censored. On average, 0.42% (SD=1.11, range=0-4.44%) of the resting-state volumes were censored. Data were spatially smoothed using a 4-mm full-width-at-half-maximum (FWHM) isotropic Gaussian kernel *(3dmerge)* and signal intensity was normalized by the mean of each voxel. EPI masks were created that included only voxels in the brain that were not excluded due to instability by *3dAutomask.* Bandpass filtering (0.01-0.1 Hz), motion censoring, and nuisance time series (estimates of motion parameters and their derivatives) were detrended from each voxel simultaneously *(3dDeconvolve, 3dTproject)* to yield a residual time series used in connectivity analyses.

Seed-based resting-state fMRI connectivity was used to determine subject-specific stimulation locations used in the subsequent concurrent TMS-fMRI experimental sessions. For each subject, a 2 mm seed in the left hippocampus (MNI: −30 −18 −18) was used in a seed-based functional connectivity analysis *(3dTcorr1D)* to identify a left lateral parietal cortex location with robust fMRI connectivity to the left hippocampal seed (mean z(r)=0.38, SD=0.05; average MNI: −53 −41 27). This was the stimulation location used for hippocampal-network-targeted (HNT) stimulation (Fig. 1C). The control out-of-network stimulation location was anatomically determined in the left supplementary motor area (SMA; average MNI: −36 −3 67), a region outside of the targeted hippocampal network. Both the HNT and SMA locations allowed the TMS coil to be positioned in the scanner without blocking the subjects’ view of the screen. Due to coil displacement during scanning, the actual achieved stimulation locations deviated from these intended targets (see below and Fig. 1C).

#### Baseline stimulation intensity determination

TMS was delivered with a MagPro X100 stimulator using a MagPro MRi-B91 air-cooled butterfly coil and MRI-compatible TMS setup (MagVenture A/S, Farum, Denmark). Resting motor threshold (RMT) was found during the baseline session in order to determine the stimulation intensity used during the experimental sessions (see below). Subjects sat at the entrance of the MRI bore with their arms resting comfortably during RMT determination. The MRi-B91 TMS coil was used to determine RMT as the minimum percentage of stimulator output (% SO) necessary to generate a visible contraction of the right thumb *(abductor pollicis brevis)* for five out of ten consecutive single pulses. Pulses were biphasic, as were pulses delivered during experimental sessions. RMT values ranged between 45.0-85.0% SO (mean=61.6, SD=10.8).

### Experimental Sessions

Following the baseline session, subjects returned for two experimental TMS/fMRI sessions on separate days to complete a 2×2 crossover design. One stimulation pattern (TBS or beta) was used during each session. Within each session, there were two study phases that differed in stimulation location (HNT or SMA). The order of these conditions (TBS or beta session; HNT-then-SMA or SMA-then-HNT within each session) was counterbalanced across subjects. Memory for scenes encoded during both study phases was tested at the end of the experimental session, after subjects finished MRI scanning.

#### Experiment Design

There were two study phases during each of the two experimental sessions. Each study phase lasted ~70 min and comprised 144 trials (288 trials total for the session). Each trial began with a white fixation cross presented in the center of the screen, during which ~ 2 s stimulation was delivered (see TMS/fMRI acquisition methods for exact timing). Immediately following stimulation, a visual stimulus was presented for 2 s. The study item was followed by a white fixation cross that remained on the screen until the next trial for a randomly varied duration between 11-19.5 s. Different visual stimuli were presented during each study phase. Complex visual scenes (50% of trials; 144 scenes total for the session) were randomly intermixed with numeric stimuli (50% of trials; 144 numbers total for the session). During the scene presentation, subjects were instructed to imagine visiting the depicted location and to rate via button press whether they would like to visit the location (right hand button) or not (left hand button). Scenes were chosen from the SUN397 dataset (Xiao et al., 2016) based on the following criteria: complex outdoor natural scenes (e.g., mountains, beaches, forests, waterfalls, deserts) without prominent humans, animals, or man-made objects; color image; image did not include text. Subjects were told that memory would be tested for all scenes following the study phase (i.e., intentional encoding). Numeric stimuli were randomly selected from the integers 1-864 and presented in white font for 2 s. Subjects used a button response to indicate if the number was even (right hand button) or odd (left hand button). Visual scene stimuli were randomly assigned to either a study trial stimulation condition or to serve as a lure during memory testing (see below) for each subject. Stimuli were presented in the center of an MRI-compatible LCD screen (Nordic Neuro Lab, Bergen, Norway) positioned at the subjects’ feet, on a gray background, viewed via a mirror attached to the head coil. Responses were made with hand-held fiber optic button boxes (Current Designs, Inc., Philadelphia, PA, USA). Subjects were told that they could make their responses during the white fixation cross following each stimulus and that response times were not important (i.e., self-paced responses).

Stimulation was delivered for ~2 s (see TMS/fMRI acquisition methods for exact timing) immediately preceding stimulus onset for 66% of scene and numeric trials (i.e., stimulation presence on), with no stimulation for the remaining trials (i.e., stimulation presence off). Long inter-trial intervals (11-19.5 s) were used to reduce stimulation carry-over effects (Huang et al., 2005). During one study phase stimulation targeted the hippocampal network via left parietal cortex (HNT), and during the other study phase stimulation targeted the SMA. Subjects came out of the scanner for a break of ~10 min between study phases for TMS coil repositioning. The order of these conditions was counterbalanced across subjects.

Prior to getting in the scanner for the study phases, MRI-navigated TMS software (Localite GmbH, St. Augustin, Germany) was used to physically mark the individualized stimulation locations on the participant’s scalp. A conformable MRI-compatible marker was affixed to the scalp at the intended stimulation location (12.7 mm x 12.7 mm re-sealable plastic bag filled with yellow-mustard MRI contrast agent; Plochman, Inc., Manteno, IL, USA). The markers were used to position the TMS coil against the subject’s head in the scanner and coil location was recorded via MRI anatomical scans during each study phase (see below).

One experimental session used TBS and the other used beta TMS, administered in counterbalanced order across subjects. For both stimulation patterns, 30 TMS pulses were delivered during the 2 s prior to stimulus onset per trial, delivered at the same intensity for each subject (80% RMT). For TBS, pulses were delivered as 50 Hz triplets at 5 Hz. For beta stimulation, pulses were delivered individually at 12.5 Hz. TMS pulses were synchronized with the MRI scan and with visual stimulus onset (see below). To acclimate subjects to the stimulation protocols and to ensure that stimulation did not cause scalp/facial twitches, a train of stimulation was applied once the subject was positioned inside the scanner and the TMS coil was positioned at the targeted location before scanning in each study phase. Stimulation intensity was lowered during one or both sessions due to technical limitations for 5 subjects. On average, TBS was delivered at 78.8% RMT (SD=1.9, range=75.0-80.0) and beta stimulation was delivered at 78.5% RMT (SD=2.2, range=74.1-80.0). The experimental sessions were scheduled at least two days apart, with an average of 27 days between sessions (range=3-84 days). For 1 subject, a session was discarded due to technical difficulties and the subject returned for a third “replacement” session. The replacement session was performed with the same location and pattern order as the discarded session, but with different visual stimuli. There were no relationships between the delay from the first to second sessions and the effect of the TBS and beta stimulation conditions on recollection memory reported below, as indicated by non-significant across-subject Pearson correlations of these variables (P’s>0.14).

At the end of each experimental session, memory was tested for the scenes that were presented during both of the study phases for that session (one study phase with HNT stimulation and one with SMA stimulation). After completing both study phases, subjects rested out of the scanner for ~15 min before taking the memory test, which was not scanned. The 144 scenes presented during the study phases were presented one at a time intermixed randomly with 144 novel lures that were not presented during study phases, in randomized order. Subjects were instructed to respond with (i) “Remember” if they specifically recalled details about the scene, (ii) “Familiar” if they recognized the scene but could not recollect details, and (iii) “New” if the scene was a lure (Yonelinas, 2002; Eichenbaum et al., 2007). Assessment of remember versus familiar responses was a key aspect of the study design, as recollection is highly related to increased hippocampal activity during encoding (Paller and Wagner, 2002; Kim, 2011; Rugg et al., 2012) and our previous experiments using hippocampal network-targeted stimulation have identified increased recollection rather than nonspecific aspects of memory performance (Hebscher and Voss, 2020). Thus, we hypothesized that the effects of HNT TBS would be specific to remember responses. Trials were self paced, with the scene remaining on the screen until a response was registered. The duration of the test phases was 20.8 min on average (range=14-29, SD=4.48).

#### Simultaneous TMS/fMRI acquisition

MRI was performed during study phases using a 3T Siemens PRISMA scanner with a single-channel transmitter/receiver head coil. Fast low-angle shot (FLASH) anatomical scans were collected between study phases to localize the actual location of the TMS coil relative to markers placed on the scalp, including a T1 sagittal (50 slices; TE 2.42 ms; TR 311.0 ms; flip angle 80°; 1.0 mm in-plane resolution; 4.0 mm thick sagittal slices with 0 mm gap; 50% phase oversampling; 256×256 mm FOV; scan duration 44 sec) and a T1 oblique axial (40 slices; TE 2.42 ms; TR 249.0 ms; flip angle 80°; 1.0 mm in-plane resolution; 4.0 mm thick axial slices with 0 mm gap; 60% phase oversampling; 256×256 mm FOV; scan duration 36 sec). These anatomical scans were later used to localize the TMS coil targeting displacement (see below).

We developed two fMRI scan sequences to interface with TMS pulses for the TBS and beta-patterned stimulation conditions. Task-based functional images were acquired using a BOLD contrast sensitive gradient echo EPI pulse sequence that contained custom programmed temporal gaps interleaved between slice acquisitions. Rather than delivering stimulation during slice acquisition, which causes TMS-induced artifact that requires volumes to be discarded (Bestmann et al., 2008; Siebner et al., 2009), TMS was delivered between MRI slice acquisitions during the inserted temporal gaps. This TMS-fMRI method did not cause artifact beyond that associated with the physical presence of the TMS coil, which produces stable artifact near the coil (see Results).

For both stimulation patterns, 30 TMS pulses were delivered during the imaging volume immediately prior to visual stimulus onset for conditions that involved stimulation. Pulses were delivered over a duration of 2000 ms in the TBS condition (Fig. 2A) and 2400 ms in the beta-patterned stimulation condition (Fig. 2B), for a total of 5760 pulses aggregate over the entire experimental session. For TBS, 107-ms temporal gaps were inserted after every two EPI slices (93 ms). During this temporal gap, a 50 Hz triplet burst (pulse every 20 ms) was delivered, with one triplet burst delivered every 200 ms during such temporal gaps (665 frames; TE 20 ms; TR 2230.0 ms; 2442 Hz/pixel bandwidth; flip angle 90°; voxel resolution 3.0 mm isotropic; 22 interleaved 3.0 mm thick axial slices angled to AC-PC alignment and centered on the longitudinal axis of the temporal lobes; 50% phase oversampling in the phase-encoding direction; 192×192 mm FOV; scan duration 24.83 min; 72 stimulated trials per scan) (Fig. 2A). For beta-patterned stimulation, 34-ms temporal gaps were inserted after each slice (46 ms), in which a single TMS pulse could be delivered, such that one pulse was delivered every 80 ms during such temporal gaps (270 frames; TE 20 ms; TR 2440.0 ms; 2442 Hz/pixel bandwidth; flip angle 90°; voxel resolution 3. mm isotropic; 30 interleaved 3.0 mm thick axial slices angled to AC-PC alignment and centered on the longitudinal axis of the temporal lobes; 50% phase oversampling in the phase-encoding direction; 192×192 mm FOV; scan duration 28.38 min; 72 stimulated trials per scan) (Fig. 2B). The scans were programmed such that the last TMS pulse would occur at the end of the TR (i.e., all pulses during the final 2000 ms of the 2230 ms TR for the TBS scan and during the last 2400 ms of the 2440 ms TR of the beta-patterned stimulation scan). Phase oversampling in the phase-encoding direction was used in both scans to shift any Nyquist ghosting induced by the presence of the TMS coil outside the brain.

**Figure 2.**
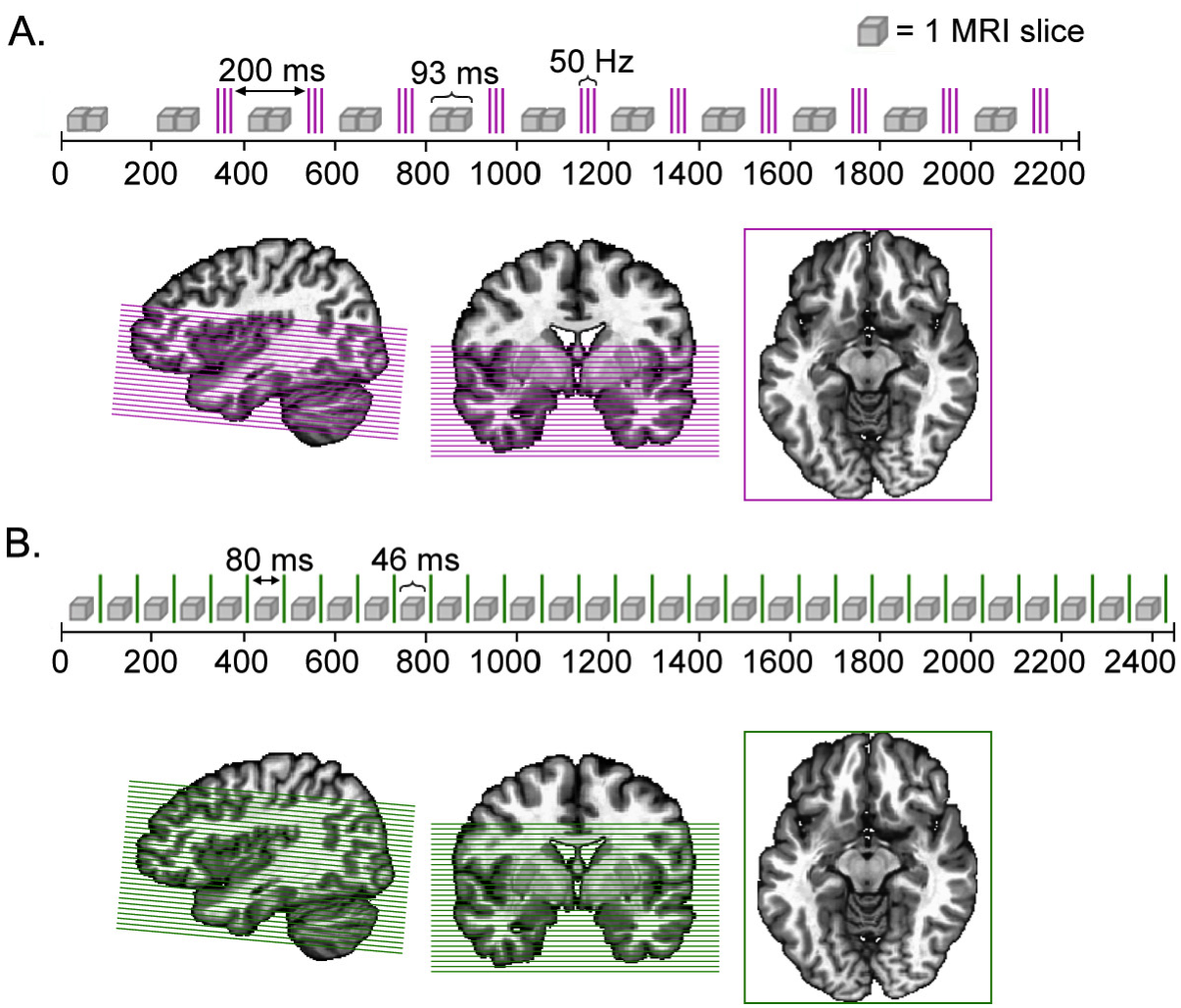
Interleaved TMS/fMRI scan sequences. Depiction of one imaging volume in the (**A**) TBS and (**B**) beta scan sequences. Each grey cube represents one MR EPI slice acquisition and each colored line indicates a TMS pulse (purple for TBS; green for beta). The extent of imaging coverage (22 EPI slices for TBS; 30 EPI slices for beta) is shown on a template brain, with EPI slices colorized to match the TMS pulses (purple for the TBS scan; green for the beta scan).

The limited coverage in both the theta- and beta-patterned task-based scans precluded whole-brain imaging but did adequately cover MTL regions of interest when the imaging volume was centered at the MTL (Fig. 2AB). Notably regions imaged directly under/around the TMS coil typically exhibit irreparable TMS-induced artifacts (Bestmann et al., 2008; Siebner et al., 2009), but the parietal cortex and SMA stimulation locations did not fall within our limited coverage. Our hypothesis-driven regions of interest were instead in downstream regions distant from the stimulation location. We confirmed image quality by subject-level visual inspection, as well as validating signal quality at the group-level (see Results). Two task-based scans using the same parameters were acquired per the two study phases during each experimental session (~70 min per phase, 144 trials per phase). Participants wore airconduction earplugs during the scans to attenuate both scanner and TMS noise.

A PC in the MRI control room received transistor-transistor logic (TTL) pulses from the MR scanner and, based on the experiment code, then sent TTL pulses to the TMS device to trigger stimulation at appropriate times. TTL pulses were sent per EPI slice acquisition to the experiment control PC, which in turn triggered the TMS device to deliver the programmed stimulation sequence. Thus, stimulation delivery was trial-specific and time-locked to the slice-based MR trigger. During the study phases, experiment events (e.g., pulse signals from the MRI and TMS, stimulator settings, participant responses, task stimuli, etc.) were monitored and recorded in output files created by Presentation (Neurobehavioral Systems, Inc. Berkeley, CA, USA), MagVenture (MagVenture A/S, Farum, Denmark), and LabChart (ADInstruments, Inc. Colorado Springs, CO, USA), as well as by the experimenter in the MRI control room. These records were used by the experimenter to update copies of raw output files with trial-specific deviations (i.e., trials were discarded if only part of the stimulation train was delivered due to coil over-heating; records reflected that the trial condition changed from “ON” to “OFF” if stimulation failed entirely during the 2-s pre-stimulus period).

#### TMS coil displacement during fMRI

We used FLASH anatomical MRI scans (see above) collected before and after the study phase fMRI scans to evaluate the actual location of the TMS coil during the experiment relative to its intended location to account for possible displacement. The scans were uploaded into the MRI-navigated TMS software (Localite GmbH, St. Augustin, Germany) and aligned to the subject’s high-resolution anatomical scan collected during the baseline session. We utilized contrast-agent markers on the TMS coil (vitamin E capsules and oil-filled tubing) and on the scalp (mustard packets, described above) to identify the position, orientation, and rotation of the TMS coil inside the scanner (i.e., 4D Matrix of coordinates) relative to the target that was identified based on resting-state fMRI for each subject (see above). The matrices were transformed with a displacement vector to estimate the cortical coordinates directly under the TMS coil. The across-participant mean MNI coordinates of the achieved HNT location was −50, −52, 32 (SD=4.8, 6.7, 6.8) for the TBS session and −49, −51, 33 (SD=4.8, 7.7, 7.3) for the beta stimulation session. The SMA location was −31, −16, 66 (SD=5.4, 9.0, 3.7) for the TBS session and −32, −13, 63 (SD=6.0, 9.3, 5.5) for the beta stimulation session. For each participant and condition, the deviation in achieved versus intended stimulation locations was calculated as the Euclidean distance. For the TBS and beta-patterned sessions, there was an average deviation of 8.47 mm (SD=4.04) for the HNT condition and a deviation of 9.02 mm (SD=4.12) for the SMA condition. The amount of deviation did not significantly vary between the two locations (P>0.7). There were no relationships between location deviation across sessions (i.e., TBS versus beta) and the differential effect of stimulation on the recollection memory measures described below, as indicated by non-significant across-subject Pearson correlations of these variables (P’s>0.36).

To confirm that the HNT and SMA conditions differentially targeted the hippocampal network as intended despite the in-scanner coil displacement, we analyzed resting-state fMRI connectivity of the achieved stimulation location (considering displacement) with the hippocampal network. Using the high-resolution resting-state fMRI scan collected at baseline, we calculated the hippocampal network as regions with robust connectivity to the left hippocampus (defined as 2-mm spherical segments centered at MNI coordinates: −23, −10, −21; −26, −14, −20; −30, −18, −18; −31 −22 −14; −30, −26, −12; see below for description of these locations in the task-based fMRI analysis). The hippocampal functional connectivity map for each subject was created by correlating (Pearson’s *r*) the spatially averaged time series of these hippocampal coordinates with every voxel’s time series (*3dTcorr*). A Fisher’s *z* transformation was applied to yield a normally distributed correlation map for each subject (*3dcalc*). Group-level voxel-wise analysis of these connectivity maps using one-sample one-tailed *t-*tests *(3dttest++)* identified clusters of contiguous voxels with robust connectivity to the hippocampal seed mask (300+ voxels with *z(r)* significantly greater than 0; t-threshold=5.2; P<0.0001). These clusters were saved as a hippocampal network mask *(3dclust;* 6,695 voxels total). This network included locations expected based on the previous literature on typical hippocampal fMRI connectivity (Kahn et al., 2008), including medial and lateral temporal, lateral parietal, ventral medial prefrontal cortex areas. For each subject, we then assessed resting-state connectivity between the achieved stimulation location to every voxel in the hippocampal network mask *(3dTcorŕ)* and spatially averaged the correlation value to obtain one overall network connectivity value for each stimulation condition for every subject *(3dmaskave).* A 2×2 repeated measures ANOVA testing the effects of stimulation location and pattern on connectivity to the hippocampal network indicated significant variation by location (F_1,12_=4.83, P=0.04, η^2^_p_=0.24), such that connectivity was significantly greater for the HNT locations (mean=0.15, SD=0.12) relative to the SMA locations (mean=0.06, SD=0.07) (Fig. 1C). Thus, differential targeting of the hippocampal network by the HNT and SMA locations was successful despite coil displacement during fMRI scanning.

To confirm that the SMA location stimulated belonged to a distinct network and therefore was a suitable out-of-network control condition, we assessed the connectivity of the achieved SMA locations using the same connectivity analysis method that was used to define the left hippocampal network but with the achieved SMA location in each subject used as the connectivity seed rather than the left hippocampus (300+ voxels with *z(r)* significantly greater than 0; t-threshold=5.2; P<0.0001; 30,275 total voxels identified). As expected, this network included canonical somatomotor regions including the majority of M1/S1, supplementary motor and somatosensory areas. The hippocampal network and SMA network had minimal overlap (7 voxels overlapping between networks out of 36,963 total voxels in both networks), confirming that the SMA location was suitable for its intended purpose as an out-of-network control.

#### Subject-level task-based fMRI processing

Anatomical scans were skull-stripped (*3dSkullStrip*) and co-registered to the Colin27 template (*auto_tlrc*). Preprocessing of the functional volumes included outlier suppression (*3dDespike*), slice timing and motion correction (*3dvolreg*), and co-registration to the anatomical scan (*align_epi_anat*). The transformations were applied simultaneously in a single resampling step *(3dAllineate).* Motion parameters were calculated for each volume as the Euclidean norm of the first difference of six motion estimates (three translation and three rotation). Volumes with excessive motion (>0.3 mm), as well as the previous volume, were flagged for censoring during the regression analyses, which is a typical threshold for task-based fMRI analysis. On average, 2.8% (SD=5.0) of the TBS HNT condition, 3.6% (SD=6.1) of the TBS SMA condition, 5.0% (SD=8.0) of the beta HNT condition, and 5.9% (SD=10.2) of the beta SMA condition time series were motion censored. There was no significant difference in the amount of censoring across conditions (all pairwise comparison *Ps*>0.10). Data were spatially smoothed using a 6-mm full-width-at-half-maximum (FWHM) isotropic Gaussian kernel *(3dmerge)* and signal intensity was normalized by the mean of each voxel. Task-based masks were created that consisted only of voxels in the brain with stable signal across the scanning sessions *(3dAutomask).*

Two general linear models (GLMs) incorporating hemodynamic response deconvolution were applied to the preprocessed data to estimate voxel-wise event-related activity regression coefficients for each trial type, separately for each stimulation condition (HNT TBS, SMA TBS, HNT beta, and SMA beta) *(3dDeconvolve).* GLMs were constructed separately per condition because differences in the scan parameters required for TBS versus beta stimulation precluded concatenation of all conditions into one GLM. In each GLM, trials were separated based on experiment condition (scenes with TMS ON, scenes with TMS OFF, numbers with TMS ON, and numbers with TMS OFF). In a second GLM, the scene trials were further sorted by subsequent memory performance (Remember, Familiar, or New responses during the test). Time points with motion spikes and time series outliers were censored. Polynomial trends and motion estimates and their derivatives were included as nuisance regressors of no interest. Condition-specific activity estimates used the duration-modulated gamma function. Each condition of interest (HNT TBS, SMA TBS, HNT beta, and SMA beta) was modeled, with each event beginning at the stimulus (scene or number) onset. Restricted Maximum Likelihood (REML) estimation methods were used to generate voxel-wise parameter estimates and measures of variability for each trial type for each stimulation condition *(3dREMLfit).* Parameter estimates from each subject were later analyzed at the group-level (see below)

### Data analysis

#### Memory performance

Performance on the scene recognition test was computed as the rate of hits (“Remember” and “Familiar” responses for studied scenes) and correct rejections (“New” response for novel lures) for each subject separately for every stimulation condition. To evaluate stimulation effects on hippocampal-dependent recollection, we calculated the proportion of hits that were recollected (“Remember” responses) for every stimulation condition. Accuracy was also calculated, as described in the Results section.

#### Group-level task-based fMRI analyses

Voxel-wise analyses were performed in order to confirm that our scan parameters provided sensitivity to expected fMRI correlates of cognitive processing (i.e., scenes but not numbers should evoke activity in parahippocampal, fusiform, and occipital regions (Stern et al., 1996; Stark and Squire, 2001) and stimulation sensations (i.e., sound emitted by stimulation should evoke activity in auditory cortex). The first contrast compared BOLD activity evoked by task stimuli (scenes versus numbers) regardless of stimulation presence, location, or pattern; and the second contrasted activation due to stimulation presence (on versus off) regardless of stimuli type or stimulation location or pattern. Subjectlevel GLMs were used to estimate voxel-wise event-related activity regression coefficients for each trial type (i.e., scenes, numbers, TMS ON, and TMS OFF) and REML estimation methods were used to generate voxel-wise parameter estimates and measures of variability for each subject *(3dDeconvolve, 3dREMLfit; see above).* GLM maps were analyzed at the group-level using generalized least squares with a local estimate of random effects variance *(3dMEMA)* to identify regions of significant difference (P<0.001, t-threshold=4.07) for scenes versus numbers and for stimulation on versus off. Data from theta-patterned and beta sessions were analyzed separately due to the differences in scan parameters.

To test whether differences in parameters between the TBS and beta stimulation scans did not significantly affect the signal quality, the Signal-to-Fluctuation-Noise Ratio (SFNR) summary value (Friedman and Glover, 2006) was assessed and compared between the two fMRI sequences. The mean signal was divided by the standard deviation of the residuals *(3dTstat, 3dcalc)* and then averaged within the limited task coverage mask over the whole session *(3dmaskave)* to yield one TSNR value per stimulation condition per subject.

To measure the effect of stimulation on activity during the memory task, performance on the retrieval task was used to back-sort fMRI data to analyze the effects of stimulation on encoding-related activity that predicted subsequent recollection (trials that were later endorsed with “Remember” responses for the main analysis, and trials later endorsed with “Familiar” and “New responses in a control analysis). Subject-level GLMs estimated voxel-wise event-related activity regression coefficients for each trial type (i.e., remembered scenes with TMS ON, remembered scenes with TMS OFF, scenes not-recollected with TMS ON, scenes not-recollected with TMS OFF, numbers with TMS ON, and numbers with TMS OFF) for each stimulation condition *(3dDeconvolve, 3dREMLfit;* see above). For scenes with “Remember” responses, there were 16 trials on average (range=6-33) per condition (trial counts did not vary by condition; Ps>0.2). Trials with remembered scenes with TMS OFF were collapsed across all study sessions to create the “combined off” condition (average number of trials=27, range=15-46). For scenes with non-recollected responses (“Familiar” and “New” responses), there were 33 trials on average per stimulation condition (range=16-64) and a mean of 17 trials in the “combined off” condition (range=8-26). Similarly, activity for numeric judgment trials was estimated, but without respect to test performance (i.e., all trials), separately for each stimulation condition, and collapsed across all study sessions for the “combined off” condition.

The influence of stimulation conditions on fMRI activity estimates for studied scenes and numeric judgment trials were tested at the group level using a region-of-interest (ROI) approach. 6-mm radius spherical regions of interest (ROIs) were defined along the hippocampal longitudinal axis in each hemisphere. The middle ROI in the left hemisphere was placed in the body of the hippocampus, centered at the coordinate that was targeted via its connectivity with parietal cortex as measured during the baseline session resting-state fMRI scan (centroid MNI coordinate: −30 −18 −18). However, because the coil was displaced, this location was not accurately targeted and was instead used simply because of its anatomical location in the body of the hippocampus. Two spheres were placed anterior to this location (centroid MNI coordinates: −23 −10 −21; −26 −14 −20), and two posterior (centroid MNI coordinates: −31 −22 −14; −30, −26, −12), in 4mm increments along the longitudinal axis. These coordinates were mirrored into the right hemisphere (centroid MNI coordinates: 23 −10 −21; 26 −24 −20; 30 −18 −18; 31 −22 −14; 30 −26 −12. The two most anterior spheres encompassed the head of the hippocampus and the middle and two posterior spheres fell in the body of the hippocampus. These spherical ROIs encompassed hippocampal as well as small portions of immediately adjacent entorhinal/perirhinal and parahippocampal cortex, which was necessary due to the relatively large voxel size. Spherical ROIs were used in lieu of more anatomically precise methods (e.g., hippocampal subfield identification) due to the limited spatial resolution imposed by the scanning parameters that are possible with the single-channel MRI head coil, which was necessary to accommodate the TMS coil. The hippocampal tail was likewise not measured, as very few voxels of this size can capture the hippocampal tail without also including substantial portions of adjacent white matter and ventricle.

The primary analysis collapsed these ROIs within each hemisphere to create separate left and right aggregate hippocampus ROIs, with the main hypothesis being that HNT TBS would increase activity evoked by scenes only in the left hippocampus. A follow-up exploratory analysis used rmANOVA to determine whether stimulation conditions had differential impact along the longitudinal axis. The goal was to identify locations along the hippocampal long axis that responded to stimulation conditions, taking into account potential functional distinctions along the anterior-posterior axis (Aggleton and Brown, 1999; Ranganath and Ritchey, 2012; Poppenk et al., 2013) as in our previous experiments investigating the variation in effects of TMS on hippocampal activity along its longitudinal axis (Wang et al., 2014; Nilakantan et al., 2019).

Voxel-wise analysis with an intentionally liberal threshold was used to evaluate whether there was any evidence for effects of stimulation in areas other than in the hippocampal ROIs, which was not expected. This analysis compared activity estimates for scenes with stimulation that were later recollected between stimulation conditions. Each subject’s voxel-wise regression coefficients for remembered scenes with TMS ON for every stimulation condition (HNT TBS, SMA TBS, HNT beta, SMA beta) was entered into repeated measures ANOVA *(3dANOVA3)* to assess voxels for a significant interaction between stimulation location and pattern. Clusters of significant interaction were identified using a liberal threshold (two-tailed P<0.05 voxel-wise threshold, F-statistic=4.54, >40 contiguous suprathreshold voxels) (*3dClust*).

### Statistical Analysis

Statistical analysis was performed using AFNI and Matlab. Group-level analysis of multiple conditions was performed using repeated-measures analysis of variance (rmANOVA), with partial eta squared (η^2^_p_) reported as the effect size. The within-subject factors corresponding to each test are provided in the Results. Greenhouse-Geisser correction was applied if the assumption of sphericity was not met (significant Mauchly’s test at P<0.05), with the corrected p-values and degrees of freedom reported (F_(GG)_) Follow-up tests avoided making all possible pairwise comparisons among the many stimulation control conditions used in the experiment by focusing on the *a priori* hypothesis that the HNT TBS condition would be unique among all stimulation conditions in that it alone would increase hippocampal fMRI activity and recollection relative to all other control conditions. HNT TBS ON was therefore the condition of interest whereas all other conditions were controls. For these follow-up comparisons, we tested whether each individual stimulation condition was significantly greater than the aggregate of all control conditions (other than the individual stimulation condition being tested) using one-tailed Students t-tests (t) or Wilcoxon signed rank tests (z) if the assumption of normality was violated (significant Shapiro-Wilk test at P<0.05), and then corrected for the number of individual stimulation conditions that were tested using the Bonferroni method. Bonferroni corrected P values are indicated as *“P_B_”* throughout the manuscript. Effect size measures were calculated as Cohen’s *d* (*d*) for the t-tests and as rank-biserial correlation (*r*) for the Wilcoxon tests. This follow-up assessment method therefore indicated whether any stimulation condition significantly increased hippocampal fMRI activity or recollection relative to the control conditions, and we hypothesized that this would only be the case for HNT TBS ON. For comprehensiveness, an exploratory analysis of all possible pairwise comparisons was used to assess whether there was evidence that any condition increased fMRI activity or recollection relative to any other condition using one-tailed Students t-tests (t) or Wilcoxon signed rank tests (z) if the assumption of normality was violated (significant Shapiro-Wilk test at P<0.05), with no correction for multiple comparisons and no reporting of effect sizes for the exploratory analysis.

### Code and Data Accessibility

Raw data are freely available on the Northwestern University Neuroimaging Data Archive (https://nunda.northwestern.edu/). Dataset identifiers will be provided with publication to permit unrestricted access to raw data. Custom code and scripts to replicate analyses will also be available via this archive.

## Results

### Validation of concurrent TMS-fMRI

The fMRI scanning methods used for TBS and beta stimulation differed in a number of critical parameters, including parameters of the scan sequence as well as the timing of scan acquisition relative to interleaved TMS pulses (Fig. 2AB). To test whether these differences affected signal quality, the Signal-to-Fluctuation-Noise Ratio (SFNR) summary value (Friedman and Glover, 2006) was calculated for each subject for the TBS and beta stimulation sessions. We expected SFNR values to be ~120, as this was the approximate SFNR value obtained when we performed the same scans on an additional subject who had the TMS coil at the same approximate locations but without any pulses delivered (average SNFR across both scans = 120.3). Furthermore, another group using the same scanner and head coil models as in this experiment reported similar SFNR values for their interleaved TMS/fMRI scans (Moisa et al., 2009). The average SFNR value for the TBS sessions was 122.5 (SD=18.3, range=94.8-160.2) and 123.4 for the beta sessions (SD=14.2, range=92.2-144.8), with no significant difference between sessions (P=0.75). Therefore, scan stability and thus, sensitivity, did not vary by scan sequence or stimulation pattern.

We performed two voxel-wise fMRI analyses to confirm expected neural activity correlates of cognitive processing within the task. That is, scenes but not numbers should evoke activity in parahippocampal, fusiform, and occipital regions (Stern et al., 1996; Stark and Squire, 2001) and stimulation sensations such as TMS sounds should evoke activity in auditory cortex. The contrast of fMRI activity evoked by scenes versus numbers, calculated across stimulation presence (on and off), and location (HNT and SMA) identified significantly greater activity (P<0.001, t-threshold=4.07) of bilateral occipital, fusiform, and posterior parahippocampal cortex as well as hippocampus for both stimulation patterns (Fig. 3A). Contrasts between TBS and beta stimulation identified no voxels with significant differences even at a liberal threshold (P<0.01 uncorrected). The contrast of stimulation “on” versus “off”, calculated across stimulation location (HNT and SMA), identified significantly greater activity for “on” (P<0.001, t-threshold=4.07) in bilateral auditory cortex for both TBS and beta stimulation (Fig. 3B). Again, the direct contrast identified no voxels with significantly different activity for TBS versus beta stimulation at a liberal threshold (P<0.01 uncorrected). Notably, although auditory-related activity due to stimulation was identified robustly, the limited imaging volume did not permit identification of likely somatosensory activation. Collectively, these analyses indicate that fMRI sensitivity was sufficient for identifying typical neural signals of scene viewing and auditory stimulation despite concurrent TMS and that there was no obvious variation in sensitivity for TBS versus beta scan parameters.

**Figure 3.**
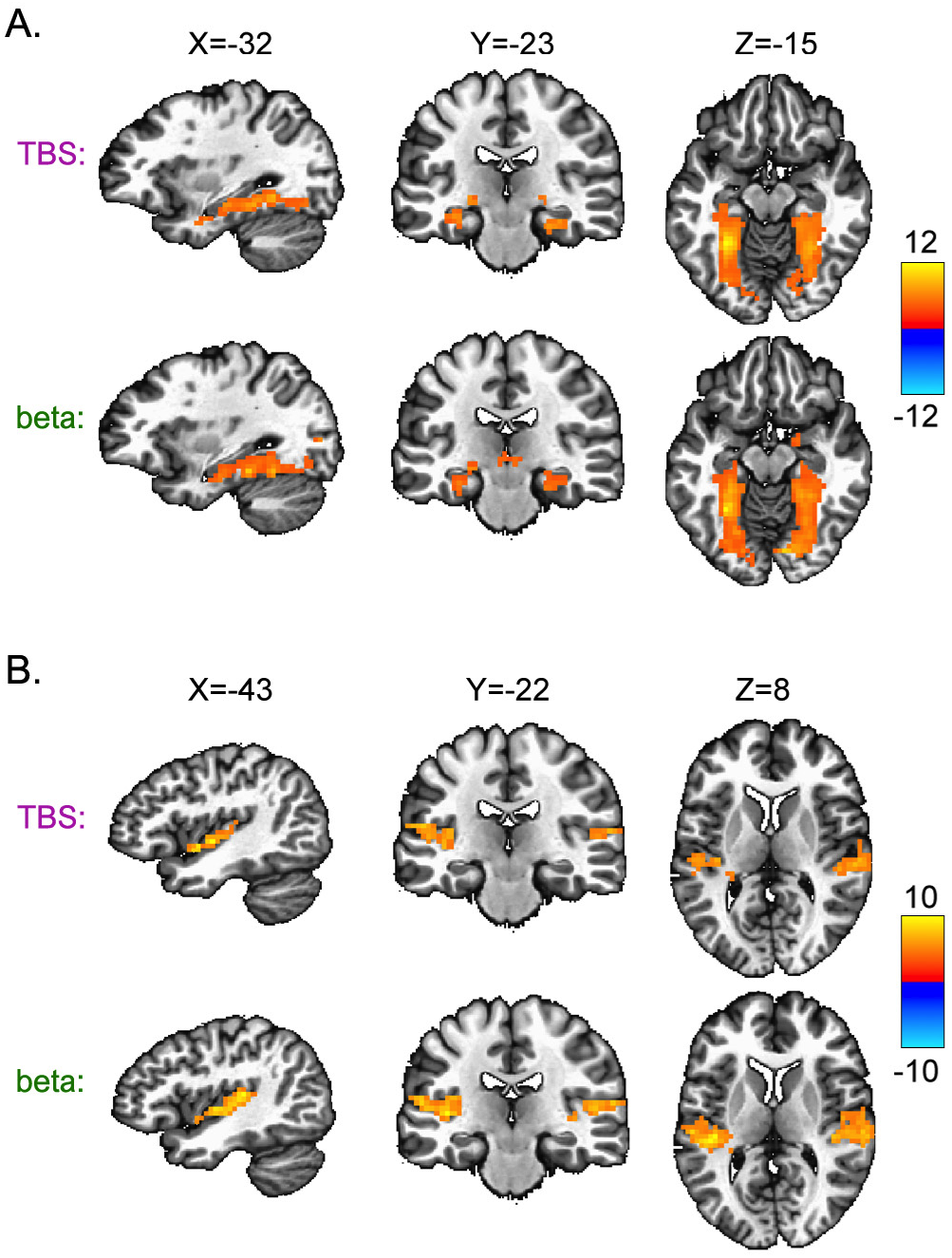
Expected fMRI signals of scene processing and stimulation sensations confirm fMRI data quality during concurrent TMS. Voxel-wise contrasts confirm that the fMRI signal could distinguish the task stimuli (scenes vs. numbers) and the presence of stimulation (ON vs. OFF). (**A**) Group-level contrast of scenes versus numbers, regardless of stimulation location or presence, identified significantly greater activation by scenes in areas that are typically scene-sensitive for both TBS and beta stimulation, including bilateral hippocampus and occipital, fusiform, and posterior parahippocampal cortex. (**B**) Group-level contrast of TMS ON versus OFF, regardless of stimulation location or the stimuli type (scenes and numbers) identified significantly greater activation for TMS ON in the bilateral auditory cortex for both TBS and beta stimulation. Direct contrasts of TBS versus beta did not identify significant differences for either comparison (see main text). Plots show supra-threshold voxels on a template brain. Color bars indicate t-statistic calculated by *3dMEMA.*

### Effects of stimulation on memory encoding

Subjects demonstrated above-chance discrimination of old/studied scenes from new scenes overall across all stimulation conditions and irrespective of whether old scenes were endorsed with “Remember” or “Familiar” responses (mean *d*’=0.81, SD=0.35; t_15_=9.15, P<0.001 vs. chance value of 0). For all stimulation conditions, the proportion of old scenes endorsed with “Remember” responses (mean=0.30, SD=0.10) was greater than the proportion of new scenes endorsed with “Remember” responses (mean=0.07, SD=0.05; t_15_=21.18, P<0.001, d=2.29). Likewise, the proportion of old scenes endorsed with “Familiar” responses (mean 0.34, SD=0.12) was greater than the proportion of new scenes endorsed with “Familiar” responses (mean=0.28, SD=0.13; t_15_=6.26, P<0.001, d=0.97). As expected, “Remember” responses were significantly more likely to be hits to old items (mean=0.84, SD=0.10) than were “Familiar” responses (mean=0.59, SD=0.10; t_15_=6.94, P<0.001, d=1.71). Thus, subjects performed the recognition task (Fig. 4A) accurately and the “Remember” responses were characteristic of high-confidence, high-accuracy recognition memory, as has been linked to the hippocampus in multiple previous experiments (Ranganath et al., 2004).

**Figure 4.**
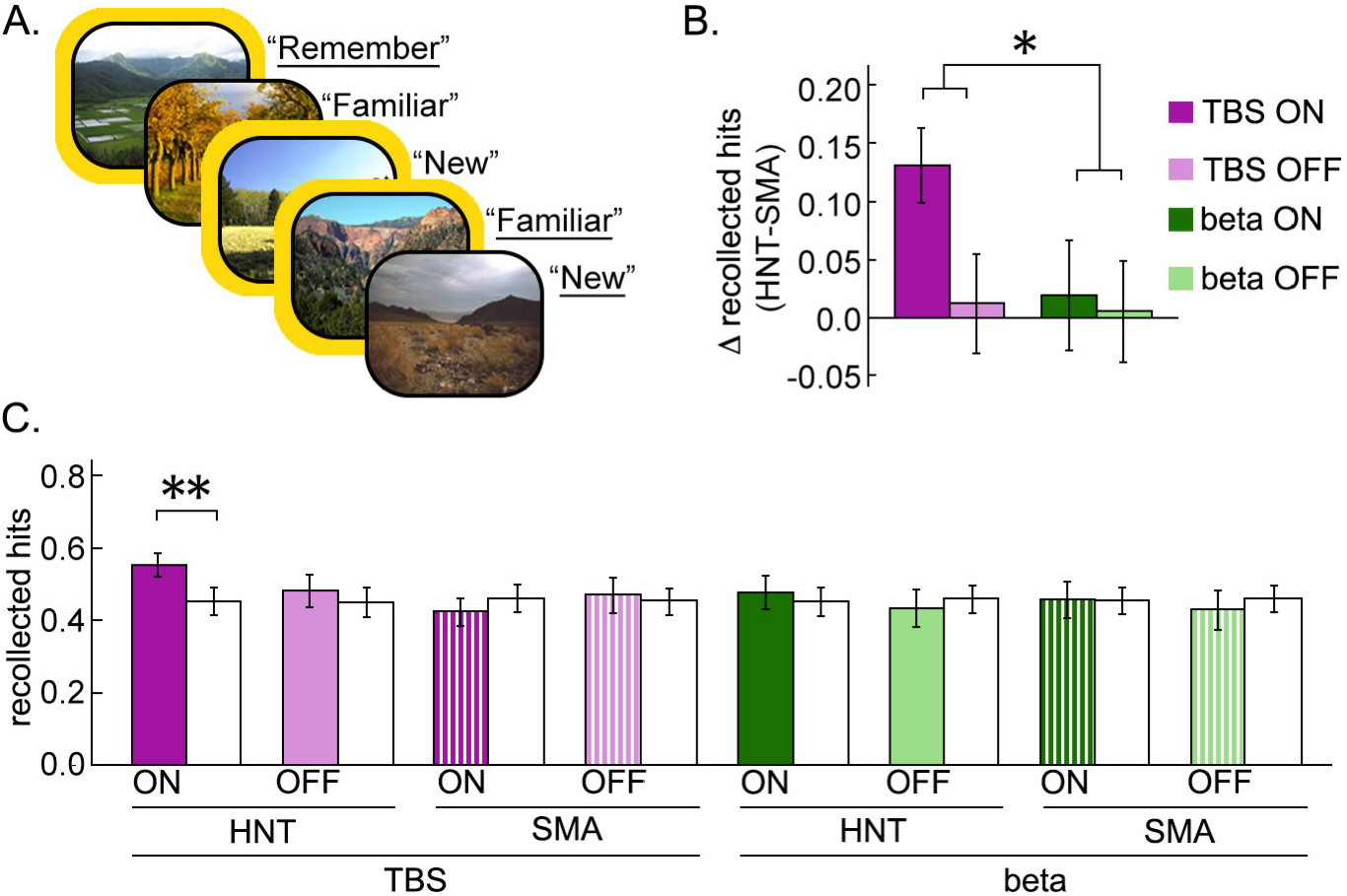
Theta-patterned stimulation of the hippocampal network during encoding selectively increased subsequent recollection. (**A**) Subjects discriminated studied scenes (old; depicted here with a yellow border) from randomly intermixed novel lures (new) using “Remember” responses to indicate the experience of hippocampal-dependent recollection, “Familiar” responses to indicate the experience of familiarity, and “New” responses to indicate lures. Example correct responses are underlined. (**B**) To identify stimulation effects on memory while holding sensory entrainment qualities (e.g., somatosensory, auditory) relatively constant, we compared the difference in the proportion of recollected hits for stimulation targeting HNT versus SMA for each of the stimulation patterns (TBS, beta, and off). * P=0.02 interaction between TMS presence and rhythm reported in text. (**C**) The proportion of recollected hits for HNT TBS was greater than for the aggregate of all control conditions. ** PB=0.008 reported in text. None of the control conditions increased the proportion of recollected hits relative to the aggregate of all other control conditions (nonsignificant values reported in text). White bars next to each condition indicate the average of the control conditions that the corresponding condition of interest (colorized bars) was tested against. Bars and error bars indicate mean ± s.e.m.

Our *a priori* hypothesis was that HNT TBS during encoding would increase subsequent recollection of scenes, as quantified as the proportion of hits endorsed with “Remember” responses, relative to all control stimulation conditions. Because of their different rhythms, TBS and beta could produce differential effects on sensory (auditory and somatosensory) entrainment of brain activity unrelated to the hypothesized effects of stimulation on hippocampal activity, and they were furthermore administered during different experimental sessions/days (Fig. 1). To test for their differential effects on recollection while accounting for these nonspecific differences, we compared the effects of TBS when applied targeting the hippocampal network (HNT) relative to the control SMA location versus the effects of beta when applied targeting HNT relative to SMA. We submitted the HNT minus SMA difference scores for the proportion of recollected hits to 2×2 rmANOVA with factors TMS presence (on, off) and rhythm (TBS, beta). There was a main effect of TMS presence (F_1,15_=5.13, P=0.04, η^2^_p_=0.23) qualified by significant interaction between TMS presence and rhythm (F_1,15_=6.53, P=0.02, η^2^_p_=0.31). As indicated in Figure 4B, TBS increased recollection when applied to HNT versus SMA to a greater extent than did beta, which had almost no differential effect when applied to HNT versus SMA. Essentially the same pattern was evident when performing this comparison without subtracting the HNT and SMA conditions. Recollection varied significantly by TMS presence (on versus off), rhythm (TBS versus beta), and location (HNT versus SMA) (3-way rmANOVA interaction F_1,11_=6.63, P=0.02, η^2^_p_=0.44). Thus, TBS and beta rhythms had different impacts on recollection when applied targeting the hippocampal network versus the control network.

The follow-up test followed our *a priori* hypothesis that HNT TBS would be the only stimulation condition to increase recollection relative to other conditions. We tested whether the proportion of recollection hits was greater for HNT TBS versus the average of all control conditions (HNT TBS OFF, SMA TBS ON, SMA TBS OFF, HNT BETA ON, HNT BETA OFF, SMA BETA ON, SMA BETA OFF), and, for comparison, tested whether recollection hits were significantly greater for each of the control conditions versus the average of all other control conditions. Consistent with our hypothesis, HNT TBS was the only stimulation condition with significantly greater recollection relative to other conditions (t_15_=3.59, P=0.001, P_B_<0.01 corrected for 8 tests, d=0.90; P_B_ = 1 for each other control condition versus the average of its respective other controls) (Fig. 4C).

An exploratory analysis of all possible pairwise comparisons among stimulation conditions was also performed, although the number of comparisons is high (56 total) and this analysis does not account for possible nonspecific differences among conditions that are relatively controlled in the primary analysis. HNT TBS was the only condition for which recollection was significantly greater than all of the other individual conditions (uncorrected P = 0.007 versus HNT TBS OFF, P = 0.03 versus HNT beta ON, P = 0.008 versus HNT beta OFF, P = 0.0005 versus SMA TBS ON, P = 0.03 versus SMA TBS OFF, P = 0.03 versus SMA beta ON, and P = 0.02 versus SMA beta OFF). In contrast, no control condition was associated with greater recollection than any one other condition (range of all uncorrected P-values = 0.08-0.99).

The experiment design used trial-specific stimulation with randomized condition order and long inter-trial intervals to ensure that the effects of stimulation were specific, as any effects of stimulation that outlasted the trial (i.e., due to longer-lasting neuroplasticity) would not have been condition specific. To more directly assess whether effects of HNT TBS on recollection outlasted the trial during which it was applied, we tested whether recollection of scenes with no stimulation (OFF) varied based on whether the scene immediately followed a trial with HNT TBS (ON) versus a trial from the same study session with no stimulation (OFF). There was no difference in the proportion of recollected hits for these two categories of non-stimulated trials (t_15_=0.66, P=0.52; two-tailed paired t-test). Thus, there was no evidence that HNT TBS continued to affect recollection on the trial after which it was applied.

Although our hypotheses concerned the effects of stimulation on recollection, we also examined effects on overall hit rates. Overall hit rate (old items endorsed with “Remember” and “Familiar” responses) did not vary by stimulation pattern, location, or presence (all 3-way rmANOVA main effects and interactions P-values>0.1) (Table 1). Notably however, although effects of stimulation on hit rates were not identified, overall memory strength could have been affected by stimulation, which could influence both the hit rate and the false alarm rate. The format of the experiment precluded calculation of false alarm rates separately for each stimulation condition because study phases for both stimulation locations (HNT and SMA) were tested at the end of each experimental session using the same set of novel lures (see Fig. 1D). Therefore, lures could be segregated based on stimulation pattern (TBS versus beta) but not based on stimulation location. The false alarm rate was significantly lower (t_15_=2.54, P=0.02, d=0.64) for TBS (mean=0.31, SD=0.17) than beta stimulation (mean=0.38, SD=0.12). Likewise, the accuracy of “Remember” responses (proportion made to old items) was significantly greater for TBS (mean=0.88, SD=0.09) than for beta (mean=0.81, SD=0.13; t_15_=2.44, P=0.03, d=0.62; two-tailed paired t-test), whereas the accuracy of “Familiar” responses did not differ significantly for TBS (mean=0.61, SD=0.11) compared to beta (mean=0.57, SD=0.13; t_15_=1.89, P=0.08; two-tailed paired t-test). Although we could not test for differences based on stimulation location, these findings indicate that TBS increased overall memory strength and recollection accuracy to a greater extent than beta stimulation, which is consistent with the primary findings that recollection was improved selectively for HNT TBS.

**Table 1.** Memory performance. For the test phase in both sessions, the total hit rate (i.e., both Remember and Familiar responses) and the proportion of hits that were recollected (i.e., Remember responses; R if hit) are provided for the scenes that were presented during the study phases for the HNT and SMA location conditions, for trials with stimulation (TMS ON) and without (TMS OFF). The total false alarm rate (i.e., both Remember and Familiar responses) and the proportion of false alarms that were endorsed with recollection (R if false alarm) are provided for scenes that were not presented during the study phase (novel foils). Statistical comparisons of interest are described in the text. Values are given as across-participant means (SD).

|  | <u>TBS session</u> |  | <u>Beta stimulation session</u> |  |
| --- | --- | --- | --- | --- |
|  | HNT | SMA | HNT | SMA |
| Total hit rate, TMS ON | 0.66 (0.10) | 0.61 (0.16) | 0.67 (0.12) | 0.66 (0.16) |
| Total hit rate, TMS OFF | 0.63 (0.13) | 0.62 (0.19) | 0.62 (0.15) | 0.64 (0.13) |
| Recollection hits, TMS ON | 0.56 (0.13) | 0.42 (0.15) | 0.48 (0.19) | 0.46 (0.21) |
| Recollection hits, TMS OFF | 0.48 (0.18) | 0.47 (0.20) | 0.44 (0.21) | 0.43 (0.22) |
| Total false alarm rate | 0.31 (0.17) |  | 0.38 (0.12) |  |
| Recollection false alarms | 0.17 (0.10) |  | 0.21 (0.23) |  |

### Effects of stimulation on recollection-related hippocampal activity during encoding

As outlined above, we hypothesized that HNT TBS would be unique among stimulation conditions in increasing hippocampal fMRI activity evoked by later-recollected scenes, and that this would occur selectively for the left hippocampus that was targeted based on its fMRI connectivity to the HNT stimulation location and based on the relatively lateralized projections from lateral parietal cortex to ipsilateral medial temporal lobe (Mesulam et al., 1977; Mufson and Pandya, 1984). Further, we hypothesized that these effects would be specific to scenes, with no impact of stimulation conditions on activity evoked by numeric-judgment trials that were randomly intermixed with scene trials during the encoding periods.

We first evaluated the effects of stimulation on activity evoked by later-recollected scenes for the left hippocampus (Fig. 5A). Activity varied significantly by stimulation condition (F_4,75_=3.40, P=0.01, η^2^=0.18, 1×5 rmANOVA for the HNT TBS, SMA TBS, HNT beta, SMA beta, and “off” conditions). The right hippocampus showed a numerically similar trend of increased activity for HNT TBS, but activity did not vary significantly by stimulation condition for right hippocampus (F_(GG)2.3,42.2_=0.76, P=0.49).

**Figure 5.**
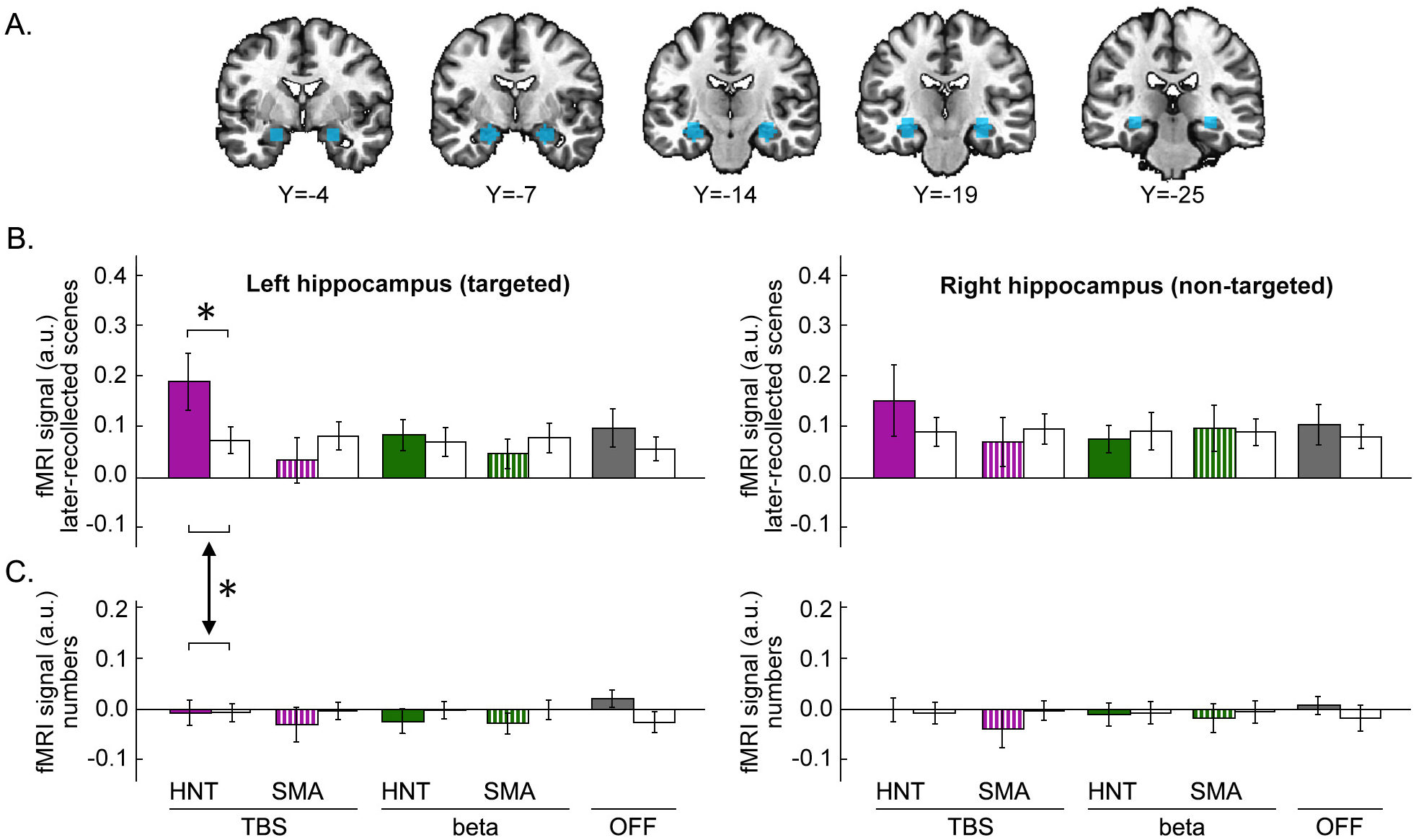
HNT TBS selectively increased left hippocampal activity during recollection memory formation. (**A**) Stimulation-evoked activity was measured within hippocampal/MTL units centered along the hippocampal longitudinal axis in each hemisphere, shown as colorized voxels in coronal slices of a template brain. The left hippocampus/MTL was targeted indirectly via its network functional connectivity (Materials and Methods) and the right hippocampus/MTL served as a non-targeted control. (**B**) fMRI activity evoked by later-recollected scenes for each stimulation condition (colorized bars) and the aggregate of all other control conditions (white bards adjacent to each colorized bars), plotted separately for left and right hippocampus. Bars and error bars indicate mean ± s.e.m. estimated fMRI activity (arbitrary units). Activity evoked by later-recollected scenes for HNT TBS was greater than for the aggregate of all control conditions. * P_B_=0.03 reported in the text. None of the control conditions increased active relative to the aggregate of all other control conditions (nonsignificant values reported in text). (**C**) Activity evoked by numbers displayed as in panel B. Significant difference of the effect of HNT TBS relative to the average of all controls for scene-evoked recollection-related activity versus activity evoked by numeric judgments is indicated (*P=0.01 as reported in the text).

The follow-up test of left hippocampal activity followed our *a priori* hypothesis that HNT TBS would be the only stimulation condition to increase activity relative to other stimulation conditions. We tested whether activity evoked by later-recollection scenes was greater for HNT TBS versus the average of all control conditions (SMA TBS, HNT beta, SMA beta, and off), and also tested whether activity was significantly greater for each of the control conditions versus the average of all other control conditions. Consistent with our hypothesis, HNT TBS was the only stimulation condition with significantly greater activity relative to control conditions (z_15_=2.61, P=0.005, PB=0.03 corrected for 5 tests; r=0.65; P_B_ for each control condition relative to the average of its respective other controls range = 0.67-1; Fig. 5B).

An exploratory analysis of all possible pairwise comparisons among stimulation conditions was also performed. HNT TBS was the only condition for which left hippocampal activity evoked by later-recollected scenes was significantly greater than for all of the other individual conditions (uncorrected P = 0.03 versus HNT beta, P = 0.002 versus SMA TBS, P = 0.02 versus SMA beta, and P = 0.03 versus combined OFF). In contrast, no control condition was associated with greater activity than any other one condition (range of uncorrected P-values = 0.07-0.93).

As shown in Fig. 5C, numeric judgments did not evoke robust activity in left or right hippocampus, relative to activity evoked by scenes (Fig. 5B), as anticipated. This activity did not vary significantly by stimulation condition for either left (F_4,75_=1.04, P>0.4) or right (F_4,75_=0.81, P>0.5) hippocampus. Focusing specifically on the HNT TBS condition of interest and left hippocampus, the effect of stimulation relative to the average of all control conditions was significantly greater for scene-evoked recollection-related activity (difference between HNT TBS bar and adjacent white bar in Fig. 5B) than it was for activity evoked by numeric judgments (difference between HNT TBS bar and adjacent white bar in Fig. 5C) (z_15_=2.30, P=0.01, r=0.58), indicating the impact of HNT TBS on activity evoked by later-recollected scenes was greater than the impact of HNT TBS on activity evoked by numeric judgments. Collectively, these findings indicate that HNT TBS selectively increased left hippocampal activity evoked by later-recollected scenes during encoding.

### Exploratory analysis of variation in stimulation effects along the hippocampal longitudinal axis

Based on evidence for anatomical and functional specialization along the hippocampal longitudinal axis (Ranganath et al., 2004; Poppenk et al., 2013) as well as our own previous evidence that network-targeted TMS can affect fMRI activity of specific hippocampal long-axis segments (Wang et al., 2014; Nilakantan et al., 2019), we performed an exploratory analysis to test for variation in the effects of HNT TBS along the longitudinal axis using five segments extending from anterior head to posterior body (Fig. 6A) (described in Methods). For left hippocampal segments, activity varied significantly by the five stimulation conditions for the two most anterior segments (F_(GG)1.9,36.0_=4.14, P=0.03, η^2^_p_=0.28; F_(GG)1.7,32.4_=3.80, P=0.04, η^2^_p_=0.25; all other Ps>0.1) (Fig. 6B). Follow-up pairwise comparisons were made among conditions using averaged activity of these two anterior segments, not corrected for multiple comparisons in this exploratory analysis. Activity evoked by later-recollected scenes was greater for HNT TBS relative to all control conditions (range of uncorrected P-values = 0.00009-0.03). Activity was also significantly greater for OFF relative to SMA TBS (t_15_=2.47, P=0.01, d=0.62), but no other control condition significantly differed from another (range of uncorrected P-values = 0.07-0.99).

**Figure 6.**
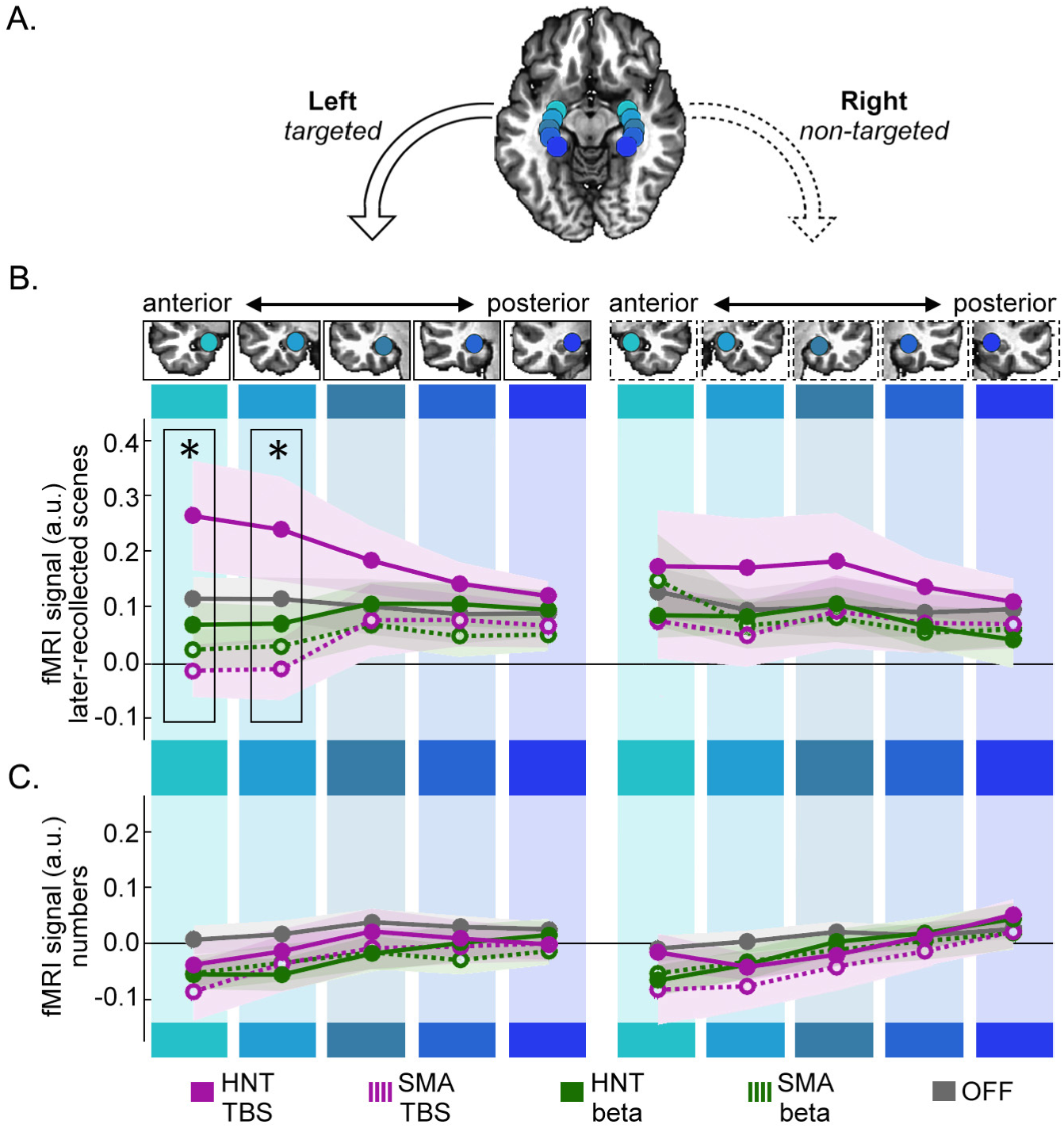
Exploratory analysis of differential stimulation effects on activity along the hippocampal longitudinal axis. The same fMRI analyses as described in Figure 5 are presented but segregated for longitudinal segments of the hippocampus in each hemisphere. (**A**) The five segments along the longitudinal axis in each hemisphere are shown as colorized spheres on a template brain. (**B**) fMRI activity evoked by scenes later-recollected was extracted from each of the spherical ROIs in the left and right hemisphere for each stimulation condition. Line plots indicate mean estimated fMRI activity (arbitrary units) ±s.e.m. (indicated by the shaded area), colorized for each of the five stimulation conditions. (**C**) fMRI activity evoked by numbers displayed as in panel B. * P<0.05 main effect of stimulation condition by five-way rmANOVA reported in text are outlined. Follow-up pairwise comparisons among stimulation conditions are described in the text.

As expected based on findings from the primary analysis (Fig. 5), the exploratory analysis performed for right (non-targeted) MTL segments yielded a numerically similar pattern of greater activity following HNT TBS relative to other conditions, but there was no significant variation by stimulation condition for any longitudinal segment (Ps>0.3) (Fig. 6B). Furthermore, as expected, MTL activity evoked by numeric judgments was minimal along the longitudinal axis and did not vary significantly by stimulation condition or segment in either the left or right hemisphere (Fig. 6C; main effect of condition and interaction of condition by longitudinal segment in the left and right hemispheres Ps>0.3). Collectively, this exploratory analysis provides preliminary evidence that effects of HNT TBS on activity evoked by later-recollected scenes were most pronounced for relatively anterior hippocampus in the targeted hemisphere.

### Voxel-wise fMRI analysis to evaluate non-hippocampal stimulation effects

To complement the ROI-based approach that was used to assess the hypothesized effects of stimulation on hippocampal activity evoked by later-recollected scenes, we performed a voxel-wise analysis with an intentionally liberal statistical threshold to determine if there was any evidence for effects of stimulation elsewhere within the FOV (Material and Methods). The rationale for using an exceptionally liberal statistical threshold is that this provides high sensitivity to the any potential effects of stimulation, with the caveat that any results would require further testing via a more rigorous approach. The voxelwise rmANOVA assessing the interaction of stimulation location (HNT, SMA) with rhythm (TBS, beta) on activity evoked by later-recalled scenes did not identify supra-threshold activity anywhere outside of the MTL. The only supra-threshold activity identified by this test was for the left anterior MTL (82-voxel cluster, peak MNI = −20, 4, −23), and this location overlaps considerably with the anterior-left hippocampal regions of interest where activity evoked by later-recollected scenes was most robustly influenced by HNT TBS (Figs. 6A). This indicates that this liberal voxel-wise test had the sensitivity to identify hypothesized effects of stimulation on activity (as it identified such effects in a similar area as in the ROI-based analysis), yet there were no robust differential effects of stimulation location and rhythm outside of the hippocampal regions of interest. The same test performed for activity evoked by numeric digits did not yield any supra-threshold clusters for either the main effects or interaction.

### No effects of stimulation on hippocampal activity during relatively unsuccessful scene encoding

In order to determine whether stimulation effects on fMRI activity evoked by scenes were specific to those scenes that were later recollected, we performed the same set of fMRI analyses previously described for the later-recollected scenes but for the scenes that were later endorsed with “Familiar” and “New” (miss) responses. As indicated above, “Familiar” responses were significantly less accurate than “Remember” responses, and “New” responses indicated memory failure. Thus, memory encoding was far less successful for this combined Familiar/New condition (44% “Familiar” responses on average, SD=6.49%) than it was for the “Remember” condition.

Stimulation condition did affect activity for this condition in the left hippocampus (F_4,75_=0.34, P>0.8) or right hippocampus (F_4,75_=1.45, P>0.2). Considering the hippocampal longitudinal axis exploratory analysis, there were also no main effects of stimulation condition or interaction of stimulation condition by the five segments for either the left or right hippocampus (main effects and interaction of condition by longitudinal segment in the left and right hemispheres Ps>0.2). The liberal whole-brain voxel-wise analysis did not find any supra-threshold clusters with an interaction of stimulation location by stimulation rhythm. Thus, the effects of HNT TBS on activity evoked by scenes were specific to the scenes that were later-recollected, with no effects of stimulation on activity for scenes comprising this low-memory condition.

## Discussion

TBS targeting the network of the left hippocampus (HNT) during encoding selectively increased fMRI activity evoked by later-recollected scenes in the targeted (left) hippocampus and increased subsequent recollection memory. These increases were rhythm- and location-specific, as they were not observed for HNT beta stimulation nor for either stimulation rhythm (TBS or beta) targeting an out-of-network control location (SMA). Further, effects were cognitively specific, as no condition (including HNT TBS) influenced hippocampal activity for scenes that were not later remembered (later-familiar and later-miss trials) nor did any condition influence hippocampal activity during numeric judgments, which do not typically evoke MTL activity (Stark and Squire, 2001). Effects were specific to the left hippocampus, which was targeted via its connectivity to the stimulated left parietal area. A numerically similar but non-significant pattern was observed for right hippocampus, which could be due to commissural connectivity and is consistent with previous findings of relatively (but not completely) lateralized effects of network-targeted TMS on hippocampal activity (Wang et al., 2014; Nilakantan et al., 2019). Finally, HNT TBS only influenced the recollective aspect of scene memory, which supports the conclusion that hippocampal function was affected, as recollection is associated with hippocampal activity during encoding (Kim, 2011; Rugg et al., 2012).

Notably, the experiment design constrains mechanistic interpretation. Stimulation was delivered as 2-sec volleys immediately before stimulus onset and with conditions intermixed across encoding trials, with intentionally long inter-stimulus intervals to avoid cumulative effects of stimulation across trials (Huang et al., 2005; Demeter et al., 2016). Any effects of HNT TBS versus control conditions must therefore have been immediate and trial specific, and indeed a control analysis provided no evidence that HNT TBS continued to affect encoding even on the subsequent trial. The experiment design and pattern of findings thus suggests that HNT TBS influenced hippocampal neural activity, as opposed to neuroplasticity and/or neuromodulatory mechanisms that can support persistent/long-lasting effects of stimulation (Cirillo et al., 2017). Notably, the design and findings also indicate that HNT TBS effects on fMRI activity were unlikely to have reflected TMS-related fMRI artefact, as different trial types were intermixed and all received the same stimulation parameters, yet effects were specific to later-recollected scenes.

The premise of the experiment was that TBS mimics endogenous theta-band neural activity characteristic of the hippocampus and hippocampal network synchrony (Buzsaki, 2002; Buzsaki and Draguhn, 2004; Lisman and Jensen, 2013) and therefore might optimally influence hippocampal memory processing via entrainment of neural activity (Thut et al., 2011a; Hanslmayr et al., 2019). Thus, if targeting hippocampal network theta activity via noninvasive TBS were successful, we expected that it would cause population synchrony of theta and therefore increase the evoked fMRI BOLD response to a stimulus that evokes processing by the affected region(s). Indeed, increased hippocampal theta activity predicts successful memory formation, particularly for complex associative information (Rutishauser et al., 2010; Fell et al., 2011; Herweg et al., 2020), including theta activity in the pre-stimulus interval (Guderian et al., 2009; Fell et al., 2011; Merkow et al., 2014) when TBS was applied in our experiment. Although our findings are consistent with this prediction, a weakness is that theta cannot be measured with fMRI and we can only infer an impact based on fMRI activity and memory performance. Further, it is also possible that recollection improvement due to HNT TBS was mediated by a mechanism other than the corresponding increase in hippocampal activity, as fMRI provides a limited window on brain activity. Full confirmation of our interpretation requires additional evidence, including direct measurement of stimulation effects on hippocampal theta.

The findings inform mechanisms by which brain stimulation influences hippocampal activity. Direct electrical stimulation of the hippocampus and its immediate entorhinal inputs via depth electrodes in human neurosurgical cases typically disrupts memory [(Coleshill et al., 2004; Jacobs et al., 2016; Goyal et al., 2018b), but see (Suthana et al., 2012; Jun et al., 2019)]. In contrast, direct/neurosurgical electrical stimulation of hippocampal network areas has yielded modest evidence for memory enhancement. Stimulation of lateral temporal cortex enhanced memory in four patients (Kucewicz et al., 2018), and closed-loop stimulation of approximately the same location caused relative enhancement compared to disruptive effects for other brain regions (Ezzyat et al., 2018). Of relevance to theta, memory enhancement was achieved in four patients receiving TBS of the fornix (Miller et al., 2015), and TBS with microstimulation of entorhinal cortex enhanced memory in several cases in which white matter (rather than gray matter) was targeted (Titiz et al., 2017). However, demonstrations of memory enhancement by invasive TBS (Miller et al., 2015; Titiz et al., 2017) did not include non-theta control conditions, and therefore do not permit strong conclusions regarding the specific role of theta.

TMS targeting the hippocampal network can generate long-lasting and robust recollection improvement and corresponding changes in hippocampal and hippocampal network activity (Hebscher and Voss, 2020), with greater aftereffects using TBS (Hermiller et al., 2019a). The current findings provide novel mechanistic insights by showing that hippocampal activity is immediately sensitive to stimulation applied to its network and mimicking its endogenous theta activity. This suggests an impact of network-targeted stimulation on hippocampal neural activity and supports the interpretation (Hebscher and Voss, 2020) that effects of noninvasive stimulation targeting this network are due to the impact of stimulation on the hippocampus.

Although sampling of activity elsewhere in the brain was limited by the fMRI methods, we found no evidence for effects of HNT TBS among locations sampled outside of the left MTL. This suggests that the hippocampus may be unique in its ability to be immediately impacted by stimulation. In contrast, previous experiments using longer stimulation trains and/or multiple days of stimulation found effects distributed throughout the network (Hebscher and Voss, 2020). Interestingly, in those studies, although stimulation was delivered at a parietal network target, the most robust effects were on fMRI activity of the hippocampus and network locations with highest hippocampal connectivity (Hebscher and Voss, 2020). It is possible that brief TBS trains affect activity in only those areas most sensitive to this stimulation pattern (hippocampus) whereas more extensive network locations are recruited with increasing stimulation.

Functionally distinct theta bands have recently been reported along the human longitudinal hippocampal axis, with slower theta (~3-5 Hz) more prevalent in the anterior and higher theta (~6-8 Hz) more prevalent in the posterior hippocampus (Lega et al., 2016; Goyal et al., 2018a). TBS was delivered at 5 Hz and the exploratory analysis suggested that it had relatively greater impact on the anterior hippocampus. This further suggests that TBS might have affected neural activity in the matching frequency band, if 5-Hz theta is more pronounced in anterior hippocampus. Formal testing of this possibility is a direction for future experiments.

The reported effects of HNT TBS are highly unlikely to have resulted from the impact of stimulation on the left parietal area where HNT stimulation was applied. This parietal location has been associated with confidence and meta-memory judgments during retrieval but not with successful memory encoding (Uncapher and Wagner, 2009; Kim, 2011). Stimulation in this experiment was applied at encoding, not during retrieval. Moreover, because effects were condition- and trial-specific, with no impact of HNT TBS even on the very next trial after it was applied, it is incredibly unlikely that stimulation had any lingering impact during the delayed test period. Because hippocampus is robustly implicated in encoding that produces later recollection whereas the stimulated area of parietal cortex is not, the experiment affords greater confidence that the effects of HNT TBS on recollection memory were related to the corresponding increases in hippocampal activity during encoding produced by HNT TBS (Tambini et al., 2018).

A limitation of the current experiment is that the memory test was given after both study phases on a given experiment session, and those study phases differed in the stimulation location (HNT versus SMA), keeping stimulation rhythm constant (TBS or beta). Thus, we could not compare effects of location on memory accuracy, as novel foils were not segregated by location. Nonetheless, we did find reduced false alarms to novel foils for TBS versus beta as well as an increase in hit rates for HNT TBS versus TBS to the SMA location. Furthermore, the accuracy of recollection, but not familiarity, responses increased for TBS versus beta. These effects on accuracy are consistent with the main findings that HNT TBS specifically impacted recollection, but future experiments will be necessary to directly test whether TBS increases accuracy only when applied to target the hippocampus versus to any location.

Hippocampal network dysfunction is related to memory impairments in many disorders (Andrews-Hanna et al., 2007; Buckner et al., 2008; Dickerson and Eichenbaum, 2010; Small et al., 2011). The current findings suggest that memory processing by the core structure of this network can be immediately and beneficially influenced via TMS of an appropriate location and rhythm. Given the immediate impact and relatively precise locus of stimulation-related activity changes, concurrent TBS with fMRI could be a powerful tool for testing hypotheses about normal hippocampal function as well as dysfunction in memory disorders.

## Acknowledgments

This research was supported by R01-MH106512 and R01-MH111790 from the National Institute of Mental Health and by T32-NS047987 and F31-NS111892 from the National Institute of Neurological Disorders and Stroke. The content is solely the responsibility of the authors and does not necessarily represent the official view of the National Institutes of Health. This research was supported in part through the computational resources and staff contributions provided for Quest, the high-performance computing facility at Northwestern University, which is jointly supported by the Office of the Provost, the Office for Research, and Northwestern University Information Technology. Neuroimaging was performed at the Northwestern University Center for Translational Imaging, supported by Northwestern University Department of Radiology. We thank Rachael A. Young and Stephen VanHaerents for contributing to data collection. We thank Plochman, Inc. for providing premium mustard.

